# MUTATIONAL PROFILE ENABLES THE IDENTIFICATION OF A HIGH RISK SUBGROUP IN MYELODYSPLASTIC SYNDROMES WITH ISOLATED TRISOMY 8

**DOI:** 10.1101/2023.01.19.524703

**Authors:** Sofía Toribio-Castelló, Sandra Castaño-Díez, Ángela Villaverde-Ramiro, Esperanza Such, Montserrat Arnan, Francesc Solé, Marina Díaz-Beyá, María Díez-Campelo, Mónica del Rey, Teresa González, Jesús María Hernández-Rivas

## Abstract

Trisomy 8 (+8) is the most frequent trisomy in myelodysplastic syndromes (MDS) and is associated to clinical heterogeneity and intermediate cytogenetic risk when found isolated. The presence of gene mutations in this group of patients and the prognostic significance has not been extensively analyzed. Targeted-deep sequencing was performed in a cohort of 79 MDS patients showing isolated +8. The more frequent mutated genes were: *TET2* (38%), *STAG2* (34.2%), *SRSF2* (29.1%) and *RUNX1* (26.6%). The mutational profile identified a high risk subgroup with mutations in *STAG2, SRSF2* and/or *RUNX1*, resulting in shorter time to acute myeloid leukemia progression (14 months while not reached in patients without these mutations, p<0.0001) and shorter overall survival (23.7 vs 46.3 months, p=0.001). Multivariate analyses revealed the presence of mutations in these genes as an independent prognostic factor in MDS showing +8i (HR: 3.1; p<0.01). Moreover, 39.5% and 15.4% of patients classified as low/intermediate risk by the IPSS-R and IPSS-M, respectively were re-stratified as high risk subgroup based on the mutational status of *STAG2, SRSF2* and *RUNX1*. Results were validated in an external cohort. In summary, the mutational profile in isolated +8 MDS patients could offer new insights for the correct management of these patients.

## INTRODUCTION

Myelodysplastic syndromes (MDS) are a group of clonal hematological disorders characterized by clinical heterogeneity and genetic diversity^1,2^. MDS display peripheral cytopenias due to ineffective hematopoiesis, morphologic dysplasia in one or more cell lineages and an increased risk of leukemia transformation. In fact, more than 30% of MDS patients progress to acute myeloid leukemia (AML). Approximately, 50% of patients show cytogenetic alterations and more than 90% have at least one genetic lesion at diagnosis^2^,^3^.

In the last decade, the identification of genetic alterations by next-generation sequencing (NGS) has changed the understanding of MDS pathogenesis^1,4^. In fact, in 2017 the presence of somatic mutations was incorporated for the first time as a new parameter for a better diagnosis of morphological subtypes^3^. Moreover, the recently published update of the diagnosis classification by both the World Health Organization (WHO) and the International Consensus Classification (ICC), defines the presence of genetic abnormalities as an important feature to consider in diagnosis of myeloid malignancies^5,6^.

The Revised International Prognostic Scoring System (IPSS-R) is the most commonly used tool for prognostic assessment of primary untreated MDS in adults. Risk-stratification remain heavily reliant on cytogenetics^7^. In fact, a cytogenetic IPSS scoring system was implemented to determine the risk of each karyotypic alteration in order to incorporate this molecular information to the IPSS-R^8^. In addition, a role for somatic mutations in the outcome of MDS patients have been described and genetic variants have been incorporated to the recently published International Molecular Prognostic Scoring System (IPSS-M). The classification of MDS patients by the IPSS-M shows improved discrimination between risk subgroups by including information from alterations in 38 recurrently mutated genes in MDS^9^.

One of the most common cytogenetic alterations found in MDS is the trisomy 8 (+8), which is present in 10% of patients with chromosomal abnormalities^10,11^. Patients with isolated +8 have an intermediate cytogenetic risk and are also characterized by clinical heterogeneity: life expectancy is different among patients, varying between several months and several years after diagnosis. Two subgroups of MDS patients with isolated +8 have been described, showing differences in overall survival (OS) based on myeloproliferative features^12^. Some of these patients showed higher number of mutations. In that way, some studies have described the presence of mutated genes in MDS displaying +8, such as *ASXL1, EZH2, STAG2, SRSF2* or *U2AF1*^12–15^. However, the role of somatic mutations is still controversial as there are no evidence of its impact in the outcome of isolated +8 in large cohorts of patients.

Given the clinical heterogeneity of MDS patients with isolated +8, accurate survival prediction and risk stratification are critical for the proper management and treatment decision-making of these patients. Therefore, the aim of the study herein was to evaluate the mutational landscape of isolated +8 and to determine whether specific somatic mutations might help to identify patients with adverse outcome in this heterogeneous group of MDS.

## PATIENTS AND METHODS

### Patients cohort and validation cohort

A total of 2 602 patients with de novo MDS were analyzed for incidence of trisomy 8. Ninety-four (3.6%) patients showed isolated +8, while 30 cases (1.1%) showed +8 in combination with additional cytogenetic alterations. Most of these combinations (80%) were complex karyotypes (CK) and, consequently, were not included in the study. Clinical and biological data are shown in Table 1. Patients with isolated +8 were compared to an IPSS-R-based matched MDS controls without +8 in a 3:1 ratio. NGS data was available in 79 out of 94 MDS with isolated +8 and in 108 out of the 282 control patients. The procedures followed were in accordance with the ethical standards of the Institutional Committee on Human Experimentation and with the Helsinki Declaration of 1975. The research was approved by the Local Ethics Committee from the University Hospital of Salamanca, Spain.

In addition, an external cohort comprising 2 494 MDS patients from the International Working Group for the Prognosis of MDS was also analyzed to validate the results^9^. A total of 94 patients showing isolated +8 were compared to 2 311 MDS cases as controls without +8, while 89 cases were excluded due to the presence of additional cytogenetic alterations together with +8 (n=81) or because of the status of chromosome 8 could not be ascertained (n=8).

### Sample procedure and DNA extraction

BM and peripheral blood (PB) samples were obtained from patients at diagnosis. Mononuclear cells were isolated by gradient density using Ficoll.

Genomic DNA was obtained using QIAamp DNA Mini Kit (Qiagen, Germany) following manufacturer’s standard protocol. The concentration of the extracted DNA was assessed by Qubit 2.0 Fluorometer system (Life Technologies, Carlsbad, CA, USA) and the integrity was analyzed by using a TapeStation 4200 (Agilent Technologies, Santa Clara, CA, USA) and a Nanodrop spectrophotometer (ND-1000, NanoDrop Technologies, Wilmington, DE, USA) by measuring absorbance ratio at 260/230 and 260/280 nm.

### Cytogenetics and FISH

Conventional G-banding analysis (CBA) data were available for all patients, and karyotypes were described in accordance with the International System for Human Cytogenetic Nomenclature^16^.

Karyotype information was confirmed by interphase fluorescence in situ hybridization (FISH) using the following commercially available probes: Vysis LSI EGR1/D5S23, D5S721 Dual Color Probe Kit, ASR; Vysis LSI D7S486 SpectrumOrange/ CEP 7 SpectrumGreen Probes, CE; Vysis D20S108 FISH Probe Kit, CE; Vysis CEP 8 SpectrumOrange; Abbott Park, Illinois, U.S.A.

### Next-generation sequencing

Samples from 187 patients underwent targeted-deep sequencing using an update of a previously validated in-house custom capture-enrichment panel^17^ (*SureSelect XT HS Target Enrichment System*, Agilent Technologies, Santa Clara, CA, USA) of 92 genes related to the pathogenesis of myeloid malignancies (Suppl. Table 1). Sequencing libraries were prepared according to manufacturer’s instructions, using unique barcodes for each sample, multiplexed and sequenced on Illumina MiSeq or NexSeq 500 sequencers.

All sequences were evaluated by using FastQC and NGSQCToolkit v2.3.3 software and aligned to the reference genome (GRCh37/hg19) using BWA v0.7.12 and GATK v3.5. A minimum quality score of Q30 was required to ensure high-quality sequencing results. Variant calling and annotation were performed using an in-house pipeline, based on the VarScan v2.3.9, SAMTools v1.3.1., and ANNOVAR bioinformatic tools. To visualize read alignments and variant calls, Integrative Genomics Viewer version 2.3.68 (IGV, Broad Institute, Cambridge, MA, USA) was used. The mean coverage of TDS was 665x (range 251-1198) and 99.5% of target regions were captured at a level higher than 100x.

For true oncogenic somatic variant calling, a severe criterion for variant filtering was applied. In addition, synonymous, noncoding variants and polymorphisms, present at a population frequency (MAF) ≥ 1% in dbSNP138, 1000G, EXAC, ESP6500 and our in-house databases, were excluded. Furthermore, those variants recurrently observed suspected of being sequencing errors were removed. The remaining variants were considered candidate somatic mutations based on the following criteria: (i) variants with ≥10 mutated reads; (ii) described in COSMIC and/or ClinVar as being cancer-associated and known hotspot mutations; and (iii) classified as deleterious and/or probably damaging by PolyPhen-2 and SIFT web-based platforms, as previously described^17^.

### Statistical analysis

Dichotomous variables were compared between different groups using the χ^2^ test and continuous variables by the Student’s t-test, Mann-Whitney and Kruskal-Wallis tests. Results were considered significant at p<0.05.

Multivariate Cox regression analysis was applied to compare clinical and molecular characteristics of patients. Survival and disease progression analyses were performed with Kaplan-Meier method and groups were compared by two-sided log-rank test. OS was defined as the period from the date of initial diagnosis to the date of death regardless of the cause. Data were censored at the last follow up in a) patients with loss of follow-up and b) patients undergoing hematopoietic stem cell transplantation (HSCT), with date of HSCT as the last point of time considered. Time to AML progression were considered as the time from the MDS to the secondary AML diagnoses and censored in patients without transformation at last follow-up or date of death.

Statistical analyses were performed using SPSS Statistics version 25 (IBM Corporation, Amonk, NY, USA), Graphpad prism version 5.03 for Windows, (Graphpad software, San Diego, CA, USA) and R version 4.0.2 with biostatistical packages (https://cran.r-project.org/).

## RESULTS

### MDS with isolated trisomy 8 have a differential mutational pattern

A total of 289 oncogenic mutations were identified in 43 genes in isolated +8 MDS patients. *TET2* was the most frequently mutated gene in isolated +8 MDS patients (38%). Interestingly, *STAG2* (34.2%), *SRSF2* (29.1%) and *RUNX1* (26.6%) were also frequently mutated in MDS patients displaying isolated +8. Nine genes were mutated in more than 10% of patients, with additional ten genes carrying mutations in 5 to 10% patients (Fig. 1a; Suppl. Table 2). Mean number of mutations per patient was 3.7 (0-10). Seventy-one (89.8%) patients had at least one mutated gene: nine patients (11.4%) had only one mutated gene while nine patients (11.4%) showed two mutated genes, and the remaining fifty-three cases (67.1%) had three or more mutated genes.

**Figure 1:**
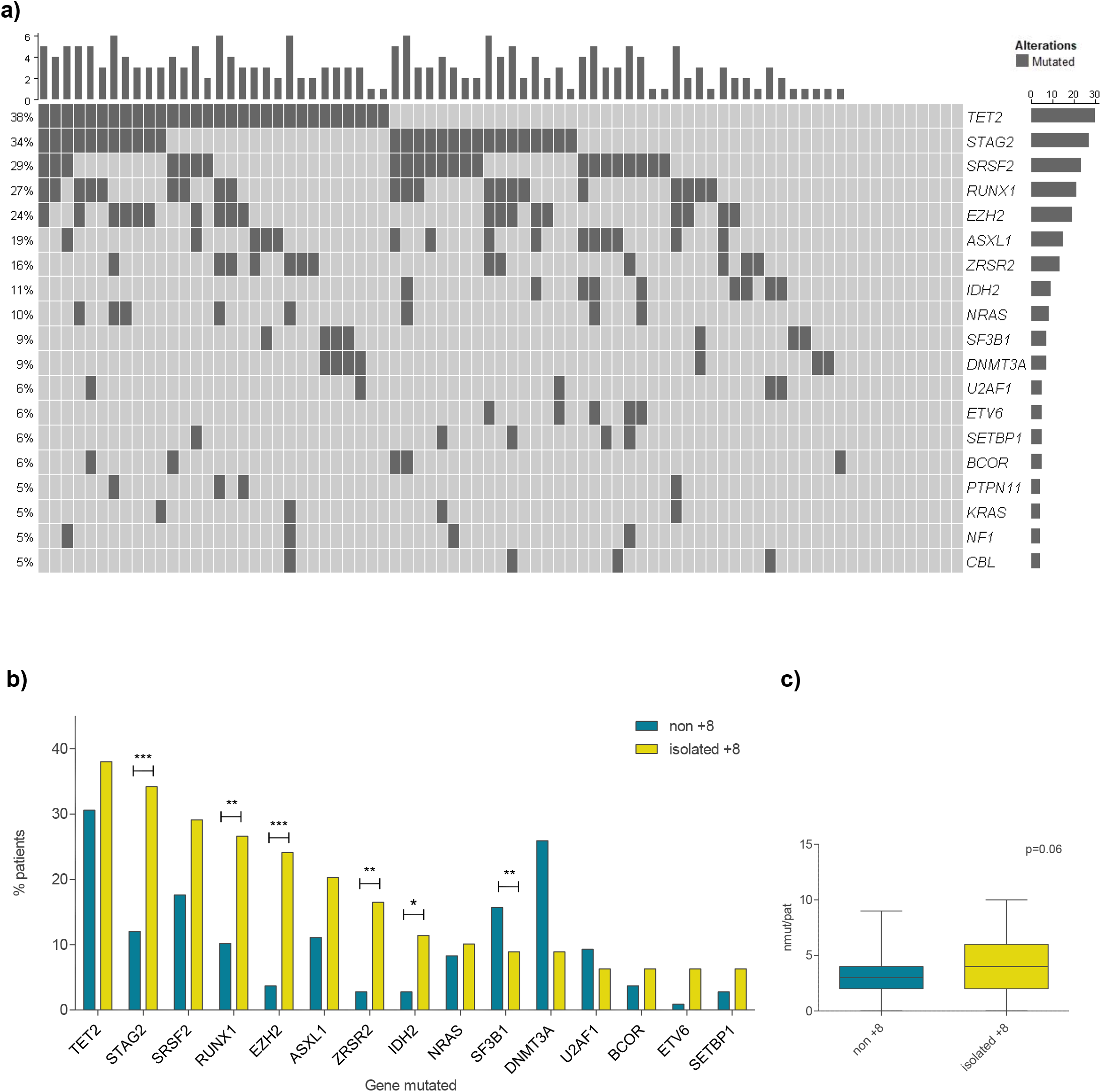
a) Mutational profile of patients with MDS and isolated +8 MDS of most frequently mutated genes (>5%); b) Comparison of top 15 most frequently mutated genes in isolated +8 and the control cohort (non-trisomy 8 MDS); c) Mean number of mutations per patient in isolated +8 compared to the non-trisomy 8 MDS group. *p<0.05; **p<0.01; ***p<0.001.

To investigate whether the pattern of recurrently mutated genes was different in this set of patients, mutational profile was compared between isolated +8 MDS and the 282 IPSS-R-matched MDS patients without +8, defined as the control group. A differential mutational pattern on isolated +8 patients was found compared to the controls (Fig. 1b). Thus, the incidence of mutations in *STAG2, RUNX1, EZH2, ZRSR2* and *IDH2* was higher in isolated +8 MDS group than in the control cohort (p<0.05). By contrast, mutations in *SF3B1* were more frequent in non-trisomy 8 MDS group (p<0.001). In addition, median number of mutations per patient was higher in isolated +8 patients compared to those MDS without +8 (4 vs 3, p<0.001; Fig.1c).

### Mutational profile allows the distinction of a worse prognosis-high risk-like subgroup in MDS with isolated trisomy 8

The presence of mutations in *STAG2, SRSF2* and *RUNX1* in isolated +8 MDS patients was significantly associated with shorter time to AML (p<0.05; Fig. 2a, Suppl. Table 3). Mutations in either *STAG2, SRSF2* and/or *RUNX1* were, therefore, considered as an entire group of alterations worsening the prognosis among patients with MDS and isolated +8 (Suppl. Table 4). In fact, isolated +8 MDS with mutations in at least one of these three genes, showed a median time to AML progression and OS similar to high/very high IPSS-R MDS patients in the control group (AML: 14 vs 11.4 months; OS: 23.7 vs 10.1 months, respectively; p>0.05; Fig.2b,c). By contrast, isolated +8 MDS patients without mutations in *STAG2, SRSF2* and *RUNX1* showed a longer time to AML progression (comparable to very low/low IPSS-R MDS patients without +8) and remained as MDS with low-intermediate IPSS-R risk in terms of OS (Fig.2b,c).

**Figure 2:**
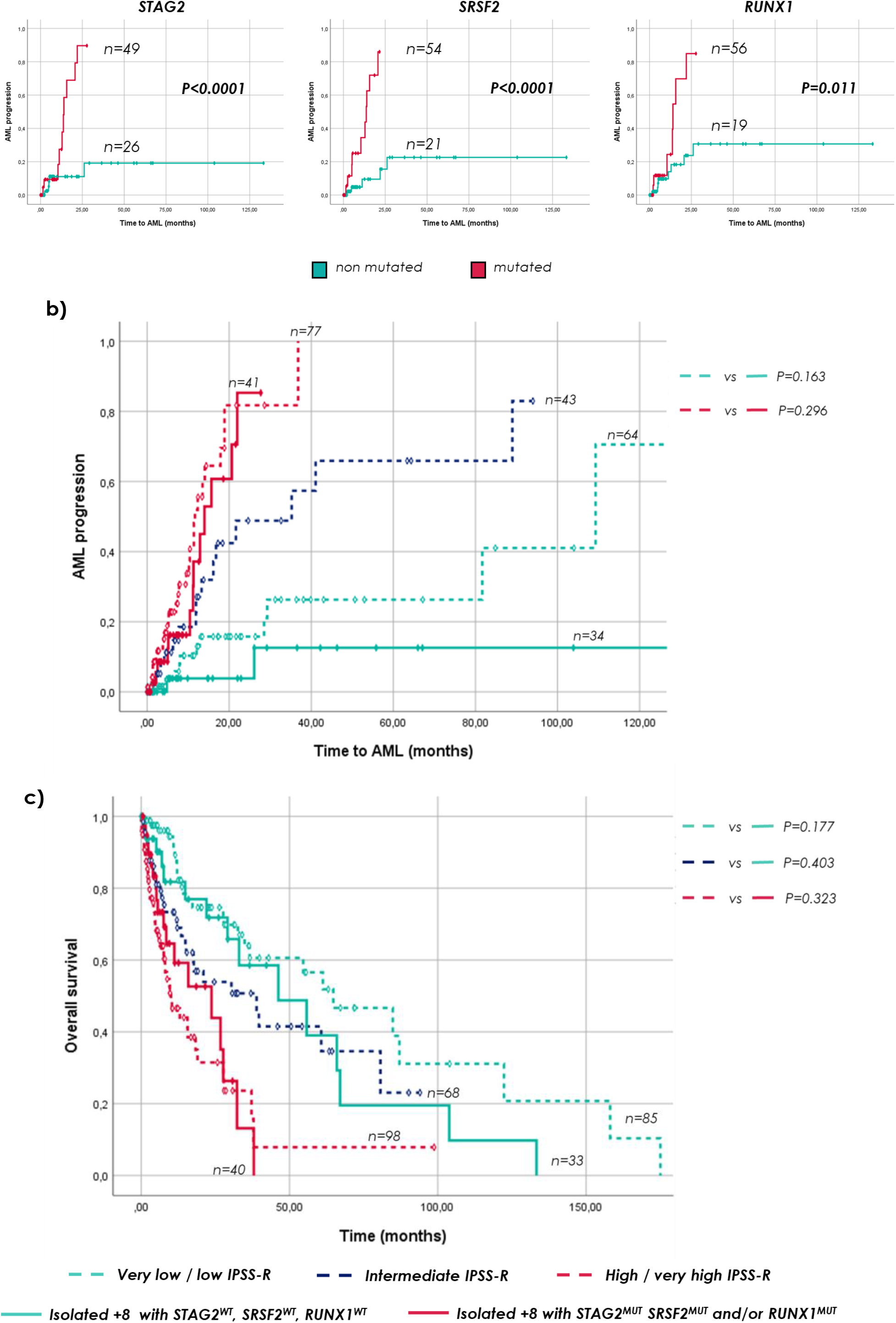
a) Time to AML progression in patients with MDS and isolated +8 according to mutational status of *STAG2, SRSF2* and *RUNX1*; b) Time to AML progression in newly described isolated +8 categories and the MDS control group stratified by IPSS-R; c) OS in newly defined isolated +8 risk categories and the MDS control cohort stratified by IPSS-R.

Moreover, time to AML progression in isolated +8 patients with mutations in *STAG2, SRSF2* and/or *RUNX1* was similar to CK (14 vs 11.7 months; p>0.05; Suppl. Fig.1a) and tended to have a comparable OS to these patients (Suppl. Fig.1b). In addition, time to AML progression in isolated +8 with no mutations in these genes was even longer than in MDS control patients with good cytogenetic risk (median not reached vs 29.3 months; p<0.05; Suppl. Fig. 1a) and similar OS was observed between them (46.3 vs 34.9 months, respectively; p>0.05; Suppl. Fig.1b).

Clinical features of patients with isolated +8 and the presence of mutations in *STAG2, SRSF2* or *RUNX1* differed from MDS patients without +8 classified as high/very high risk by the IPSS-R in levels of hemoglobin and percentage of BM blasts (Table 2). Interestingly, despite having higher hemoglobin levels and lower counts of blast in BM, no differences were found in patient’s outcome in terms of both AML progression and OS (Fig. 2, Table 2). Otherwise, patients with isolated +8 and without mutations in neither *STAG2, SRSF2* nor *RUNX1*, showed a similar age, hemoglobin level, absolute neutrophils count and outcome than MDS without +8 classified as low risk by the IPSS-R, but a higher percentage of BM blasts and lower levels of platelets and ring sideroblasts (Fig. 2, Table 3).

Consequently, the presence of mutations in *STAG2, SRSF2* and/or *RUNX1* distinguishes between two different prognostic subgroups displaying specific clinical characteristics and outcomes similar to high risk (with at least one mutation in these genes) and low risk (without any of these genes altered) MDS patients without +8.

### Mutational status of *STAG2, SRSF2* and/or *RUNX1* is an independent prognostic factor associated with shorter time to progression to AML and overall survival in MDS with isolated trisomy 8

In a multivariate Cox regression analysis, clinical parameters included in the IPSS-R were separately evaluated. In this regard, age (≥75 years), percentage of BM blasts (≥5%), number of cytopenias (>2) and mutational status of *STAG2, SRSF2* or *RUNX1* were considered. Alterations in these genes were associated with shorter time to AML (p<0.01; Fig.3a). Interestingly, the number of mutations per patients did not associate with time to AML (p>0.05; Suppl. Fig. 2b) indicating that mutations in either *STAG2, SRSF2* and/or *RUNX1* are enough to predict early disease progression in isolated +8 MDS patients.

**Figure 3:**
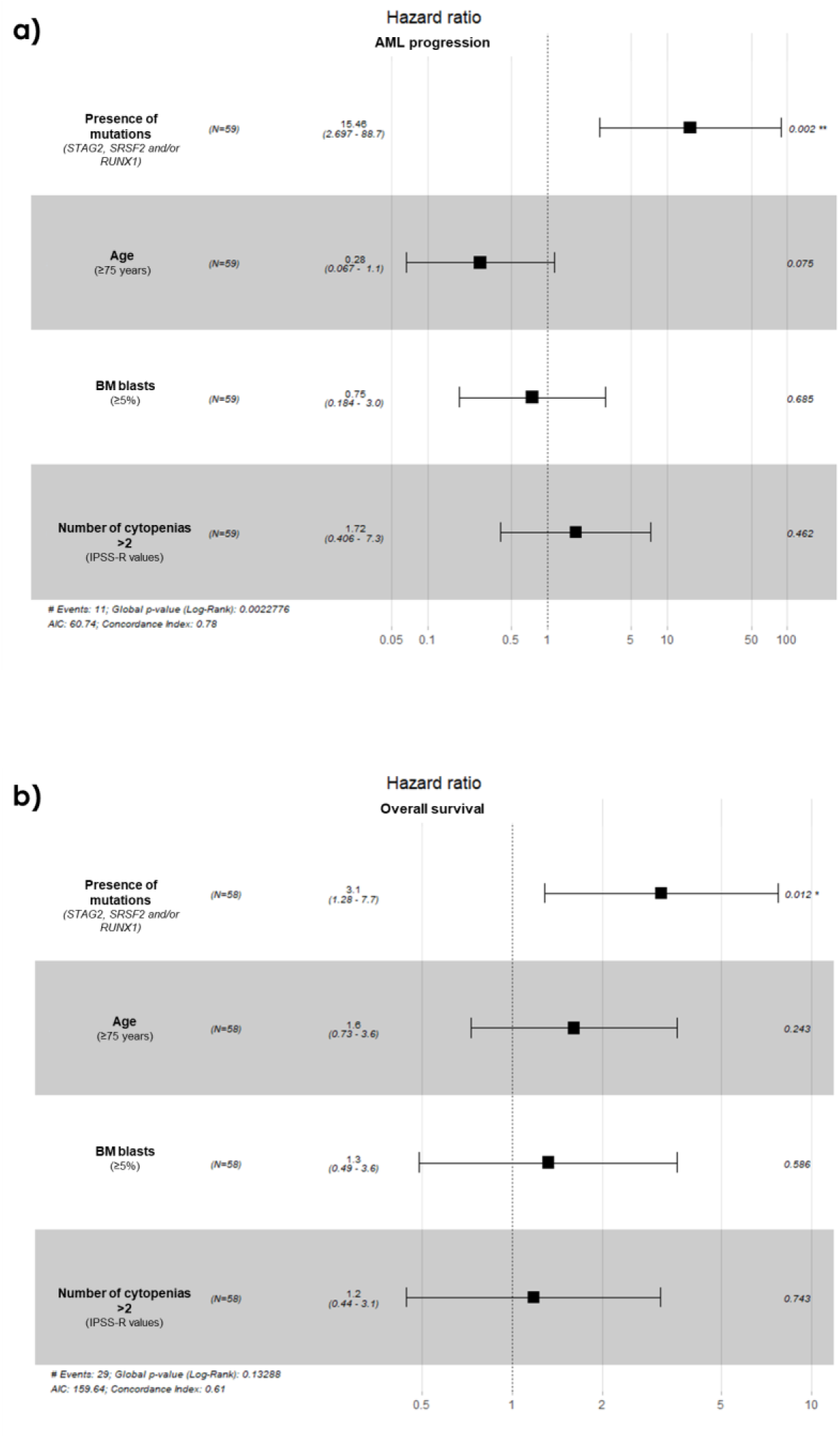
a) Multivariate analysis of time to AML progression in the isolated +8 MDS cohort; b) Multivariate analysis of OS in patients with MDS and isolated +8.

OS was also evaluated in a similar multivariate analysis considering age (≥75 years), percentage of BM blasts (≥5%), number of cytopenias (>2) and mutational status of *STAG2, SRSF2* and *RUNX1*. Only mutations in these genes remained as independent predictor of worse outcome (p=0.01; Fig.3b). By contrast, number of mutations per patient (>4) did not associated to shorter OS (Suppl. Fig. 2b). Furthermore, multivariate analyses also revealed that when considering a percentage of BM blasts higher than 10%, the mutational status of *STAG2, SRSF2* and *RUNX1* was still an independent prognostic factor of shorter OS, together with >10% blasts counts and age (Suppl. Fig. 2c,d).

### The newly defined prognostic mutations could refine prognosis of patients with isolated trisomy 8 within the IPSS-R and the IPSS-M systems

Patients with MDS and isolated +8 were classified within both IPSS-R and IPSS-M. A comparison between our defined mutational prognostic signature and both IPSS-R and IPSS-M was assessed to evaluate whether mutational status of *STAG2, SRSF2* and *RUNX1* could add value to these prognostic scoring systems in isolated +8 MDS. Remarkably, a 23.5% of patients categorized as low risk by the IPSS-R showed a high-risk mutational profile (this is, mutations in either *STAG2, SRSF2* or *RUNX1*), while a 52.4% of patients within the intermediate risk patients had a high-risk mutational profile. Overall, the risk of 39.5% low/intermediate patients were underestimated by the IPSS-R. By contrast, 16.7% and 35.3% of patients classified as very high and high risk by the IPSS-R, respectively, were included in the low risk subgroup without these mutations when analyzing mutational profile (Fig. 4a).

Otherwise, only one patient (5.6%) in the low IPSS-M and a 37.5% of patients with moderate low IPSS-M risk were classified as high risk by mutational profile (which correspond to a 15.4% of low/moderate low risk patients by the IPSS-M) (Fig. 4b). As a result, most of the patients showing mutations in *STAG2, SRSF2* and/or *RUNX1* were categorized as moderate high, high or very high risk by the IPSS-M (88.9%) while only 51.6% were categorized as high or very high by the IPSS-R. Consequently, the risk of isolated +8 patients with mutations in *STAG2, SRSF2* and/or *RUNX1* was refined in a total of 11.1% by the IPSS-M and in 48.4% of cases by the IPSS-R (Suppl. Fig. 3).

**Figure 4:**
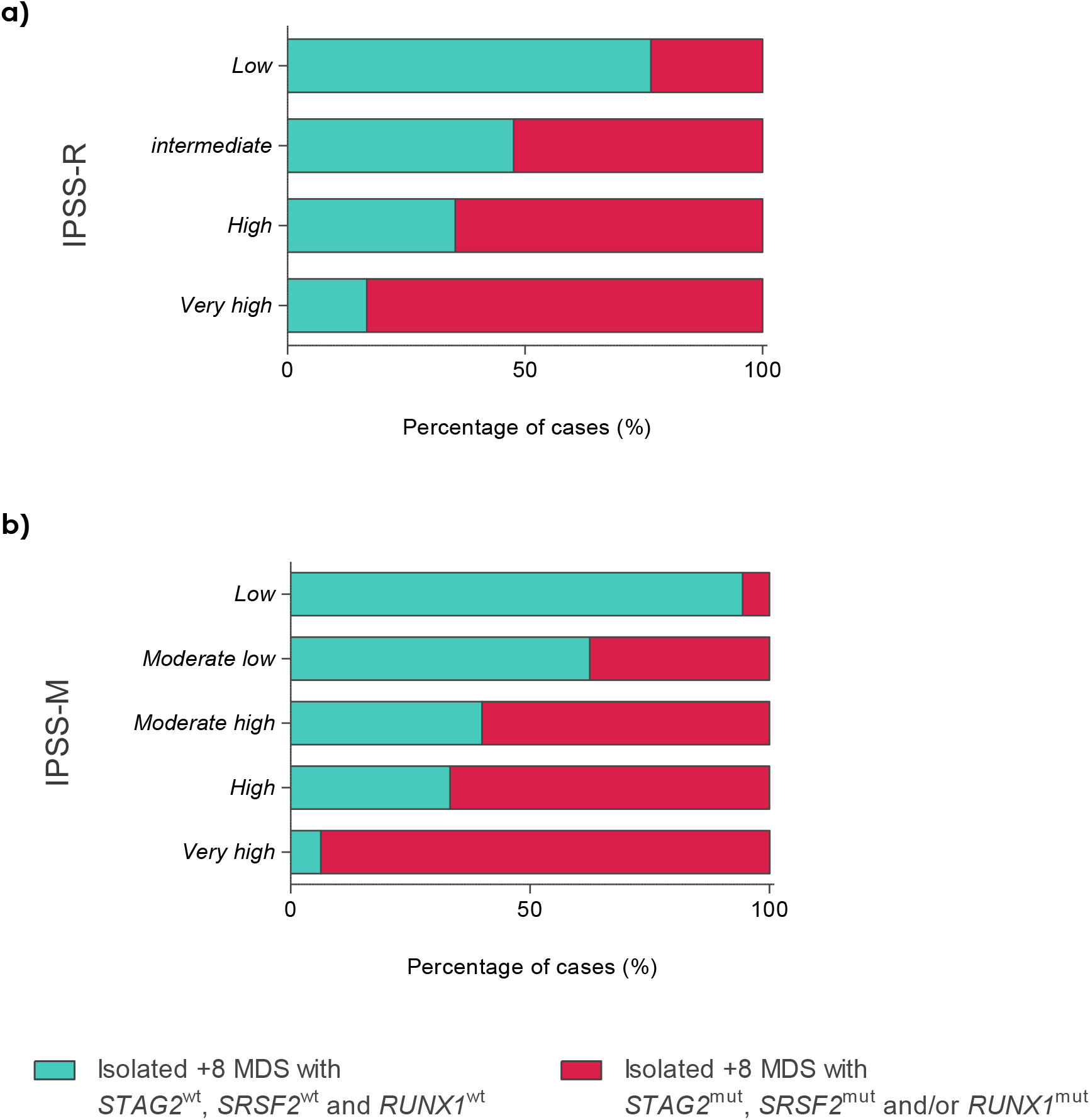
a) Percentage of patients stratified within the IPSS-R classified as low risk-like and high risk-like based on mutation status of *STAG2, SRSF2* and *RUNX1*; b) Percentage of patients classified within the IPSS-M categories regarding the risk based on mutational profile.

### *STAG2, SRSF2* and *RUNX1* differentiate a subgroup of isolated trisomy 8 MDS with worse outcome in an independent validation cohort

Finally, to further characterized the mutational landscape associated to isolated +8 in MDS, additional analyses were done in an external validation cohort. Thus, a previously published MDS population comprising 2 494 patients from the International Working Group for the Prognosis of MDS was evaluated^9^.

A total of 94 MDS patients presented isolated +8 (3.8%) and clinical characteristics were similar to those of the discovery cohort (Suppl. Table 5). In this set of patients, the most frequently mutated genes were *ASXL1* (44.7%), *TET2* (39.4%) and *STAG2* (36.2%). The mutational incidence observed in most of the genes were similar to those of our cohort of MDS with isolated +8 patients (Suppl. Fig.4). Twenty-three (24.5%) and twenty-one (22.3%) patients showed mutations in *SRSF2* and *RUNX1*, respectively. In addition, median number of mutations per patient was higher in isolated +8 patients compared to MDS without +8 (4 vs 3, respectively; p<0.0001), as in our discovery cohort.

As in our own cohort, the presence of mutations in *STAG2, SRSF2* and/or *RUNX1* were able to distinguish a worse prognosis subgroup of patients. In fact, patients with mutations in these genes showed a similar time to AML and OS than high/very high risk IPSS-R MDS patients (2.2 years of AML transformation in the high risk patients without +8 and 1.6 and 1.2 years of OS, respectively; p>0.05; Suppl. Fig.5a,b). Multivariate analysis also revealed an independent prognostic value for these mutations in isolated +8 MDS (p<0.05; Suppl. Fig. 5c).

## DISCUSSION

MDS patients are characterized by clinical and genetic heterogeneity that generally translate into varied outcomes. In fact, some cases with isolated +8 progress quickly to AML while others remain stable during years. Some studies have analyzed the clinical variability in patients with MDS and isolated +8, precluding any clear-cut conclusions^18,19^. However, most of these researches were carried out before the implementation of NGS into clinics and evidence a lack in understanding the pathogenesis of isolated +8 patients. In view of the limited evidence on the specific impact of mutational landscape in the isolated +8 MDS setting, the present study was undertaken to determine whether mutational profile might help to stratify patients with isolated +8. The deeply targeted gene sequencing reported in this work provides an unprecedented overview of genomic landscape of MDS in a context of isolated +8 and throw evidence on how genetic lesions play a role in the differential outcome of MDS patients with isolated +8.

The comparison of mutational landscape between isolated +8 and non-trisomy 8 MDS showed that patients with isolated +8 displayed a differential mutational profile characterized by a slight increase in the mean number of mutations per patient (Fig. 1c). In addition, mutations in *STAG2, RUNX1, EZH2, ZRSR2* and *IDH2* were more frequent in the isolated +8 MDS than in the control group (Fig. 1b). Remarkably, the proportion of patients with isolated +8 MDS and mutations in *STAG2* was near three times more that in control MDS as previously reported^12^. As expected, the incidence of mutations in *SF3B1* was lower in the isolated +8 cohort compared to control MDS due to these mutations have been extensively associated to MDS with ring sideroblasts^20–22^ or normal karyotypes^23^, which is concordant to the low incidence of these alterations in our cohort. By contrast, our results did not show *TP53* as frequently mutated in MDS with isolated +8, which was consistent with the high association of mutations in this gene to both complex karyotype and 5q deletion previously described in MDS^24–26^. All this data together could indicate a specific spectrum of genetic lesions associated to isolated +8.

The presence of *STAG2, SRSF2* and *RUNX1* mutations, alone or in combination, shortening both the time to leukemia transformation and the OS within MDS patients with isolated +8. Despite the well-known key prognostic significance that percentage of BM blasts plays in MDS, patients with isolated +8 and mutations in *STAG2, SRSF2* and/or *RUNX1* showed similar outcomes to high/very high risk control patients, even with significant lower levels of blasts in BM (Table 2). By contrast, higher levels of BM blasts were observed in isolated +8 MDS without these mutations, when compared to low risk MDS controls. In addition, the low frequency of *SF3B1* mutations in isolated +8 patients might explain the lower levels of ring sideroblasts and platelets (of which high levels have been associated with *SF3B1* mutants^20^) observed in these patients (Table 3). All these data suggest that not clinical, but genetic information might have an essential role in the prognosis of patients with isolated +8.

In fact, not number of mutations but alterations in these genes become an independent prognostic factor in isolated +8 MDS patients with a low percentage of blasts (<5%), as shown in the multivariate analysis in the discovery and the validation cohorts (Fig. 3, Suppl. Fig. 5c). Despite the prognostic relevance of BM blast in the outcome of MDS^19^, the presence of *STAG2, SRSF2* and/or *RUNX1* were the only independent prognostic factor in this set of patients with lower counts of BM blasts (<5%).

When IPSS-R was applied to our cohort of MDS with isolated +8, 23.5% of patients included in low risk category and 52.4% of intermediate risk cases categorized by the IPSS-R, showed worse outcomes and were described as high risk isolated +8 patients due to the presence of mutations in *STAG2, SRSF2* and/or *RUNX1* (Fig. 4a). In addition, 30.4% of patients categorized as high and very high risk by the IPSS-R were considered as low risk by mutational profile. Therefore, our results suggest that comprehensive genomic analyses might refine the risk classification in a subset of MDS patients with isolated +8.

Otherwise, these isolated +8 differentiated subgroups were validated by the new IPSS-M classification, as a high number of patients (88.9%) with mutations in *STAG2, SRSF2* and/or *RUNX1* were classified as moderate high, high and very high risk by the IPSS-M (Fig. 4b). Of note, a 37.5% of patients with moderate low risk by the IPSS-M showed a high risk based on status of *STAG2, SRSF2* and *RUNX1*. In this regard, while *SRSF2* and *RUNX1* are genes considered as relevant factors increasing the score, *STAG2* shows less relevance in the IPSS-M^9^. However, *STAG2* is the secondly most frequent mutated gene in isolated +8 MDS patients and play a role in disease progression and OS^17,28^, as has been demonstrated in this work. Therefore, the score associated to each somatic mutation might vary depending on the subset of MDS patients studied. Specifically, analysis of mutations in *STAG2, SRSF2* and/or *RUNX1*, in addition to the IPSS-M, could refine prognostic information of patients presenting isolated +8. In addition, our results could be of special interest in patients classified as moderate low by the IPSS-M, where *STAG2* mutations might be worsening the prognosis.

It is well known that the heterogeneity of hematological diseases as MDS makes difficult the management of patients. Indeed, only high risk MDS patients are eligible for receiving HSCT or being treated with HMA agents due to the increasing risk of leukemia transformation^29^. The identification of patients likely to be treated in the earliest steps of the disease would enable to anticipating to clinical deterioration of patients and increases options of stabilizing the disease in a good state of patient health. In fact, both shorter time to initiating HMA treatment and immediate HSCT at the time of diagnosis maximized OS and complete responses in high-risk MDS patients^30-32^, while delays during the first cycles of therapy adversely affect OS, independently of the IPSS-R status of the patient^33^. Consequently, identifying a subgroup of isolated +8 MDS with higher risk could transform the therapeutic approach of these patients.

Summarizing, the identification of somatic mutations has shown to be useful in distinguishing different risk subgroups in isolated +8 MDS patients. In fact, the presence of mutations in three genes (*STAG2, SRSF2* and *RUNX1*) would be enough to determine the course of the disease in terms of OS and AML progression. Overall, these data could open new horizons with the identification of drug targets and drive to personalized medicine by changes in the management in a high risk subset of patients with isolated +8 MDS.

## ACKNOWLEDGES

The research leading to these results has mainly received funding from “*Gerencia Regional de Salud de Castilla y León, GRS 2341/A/21*”. In addition, the NEMHESYS project (Grant Agreement 612639-EPP-1-2019-1-ES-EPPKA2-KA), co-funded by the Erasmus+ Programme of the European Union, has trained some of the researches involved in the work.

S.T.C. is a recipient of a predoctoral fellowship from the “*Consejería de Educación de la Junta de Castilla y León, EDU/601/2020”* and M.D.R is supported by a fellowship from the Spanish Society of Hematology (FEHH).

We are grateful to I. Rodríguez, S. González, S. Santos, C. Miguel, T. Prieto, M. Á. Ramos, A. Martín, A. Díaz, A. Simón, M. del Pozo, V. Gutiérrez and S. Pujante from Centro de Investigación del Cáncer, Salamanca, for their technical assistance.

## CONTRIBUTIONS

JMHR and TG conceptualized the study and designed research. STC, SCD, and AVR performed the research and analyzed the data. STC undertook methodology. STC and MDR wrote the paper. MDC, MDR, TG and JMHR contributed to scientific discussion, data interpretation, and paper revision. SCD, ES, MA, FS, MDB and TG provide patient samples and clinical data. MDC and JMHR supervised the study and acquired funding. All authors reviewed and provided edits and/or helpful comments on the manuscript.

## FIGURE LEGENDS

**Table 1:** Clinical and biological features of isolated +8 MDS cohort.

*MDS-SLD: MDS with single lineage dysplasia; MDS-MLD: MDS with multilineage dysplasia; MDS-RS: MDS with ring sideroblasts; MDS-del(5q): MDS with deletion of 5q; MDS-EB-1: MDS with excess of blasts type 1; MDS-EB-2: MDS with excess of blasts type 2.

**Table 2:** Clinical features of isolated +8 MDS patients with mutations in *STAG2, SRSF2* and/or *RUNX1* compared to high risk IPSS-R and high risk IPSS-R MDS controls.

**Table 3:** Clinical characteristic of patients with isolated +8 MDS without mutations in *STAG2, SRSF2* and *RUNX1* compared to low risk IPSS-R control patients.

